# Mitochondrial Dynamics Through the Lens of Nonlinear Chemical Oscillations: A Theoretical Exploration Inspired by the Belousov-Zhabotinsky Reaction

**DOI:** 10.1101/2025.05.15.654228

**Authors:** Ayalur Raghu Subbalakshmi, Tamara Mirzapoiazova, Prakash Kulkarni, Ravi Salgia

## Abstract

In biology, oscillations are observed across a wide spectrum of processes and systems. Oscillatory systems are typically leveraged to transmit information within cells. However, they can also serve to transmit information between organisms underscoring their fundamental role in regulating transitions, maintaining stability, and responding to environmental stimuli. In this study, we explore mitochondrial fission and fusion dynamics through the framework of the Belousov-Zhabotinsky (BZ) reaction, a hallmark of non-equilibrium system that exhibits periodic changes in reactant concentrations through autocatalysis and feedback regulation. We observed that mitochondrial changes followed an oscillatory dynamic where the fission, fusion and intermediary factors undergo oscillations. Also, by modelling comparison with publicly available datasets of diseased condition before and after therapy, we observed similarities, where under diseased condition there is increased concentration of the fission and fusion factors but upon treatment the concentration of the intermediary factors increase. Also, patient survival data analysis showed that increase in fission and fusion factors correlated with increased deaths but when there is increase in the intermediary factor concentration, we see better patient survival. These results highlight the possibility of targeting mitochondrial dynamics as a potential strategy for therapeutic development for diseases such as cancer where mitochondrial dynamics is dysregulated.

## Introduction

Oscillatory and cyclic processes are fundamental to the functioning of biological systems and these processes enable precise temporal regulation across biological scales. From a cellular level to an organismal level critical processes are seen to be governed by oscillatory processes that help in the maintenance of homeostasis and increase adaptability (1,2). And among these oscillatory processes cell cycle is one of the very well-studied examples. Cell cycle is a tightly regulated process where a series of events results in cell division and proliferation. Cell division is characterized by oscillatory changes in cyclin-dependent kinases (CDK), that results in periodic transitions between the cell-cycle phases (G1, S, G2, and M) (3–6). Another example of biological oscillations is circadian rhythms. Circadian rhythms regulate physiological processes over a 24-hour period, relying on feedback loops between clock genes and their protein products, namely CLOCK and BMAL1 (7–10). Other examples include calcium signaling oscillations in response to stimuli (11,12), oscillations in metabolic signaling (13), and heartbeat regulation (14,15), all of which rely on feedback, activation, and inhibition to sustain periodicity. Beyond these well-established cycles, oscillations also drive phenotypic switching, as seen in c-Myc dynamics regulating cellular states(16)and phosphorylation mediated conformational changes contributing to phenotypic heterogeneity as seen in the PAGE4 (17). These myriad systems highlight a common underlying principle that in biological systems oscillatory dynamics are essential for regulating transitions, maintaining stability, and responding to environmental stimuli.

The principles of cyclic processes in biology find a striking parallel in chemical oscillators, such as the Belousov-Zhabotinsky (BZ) reaction. The BZ reaction, a hallmark of non-equilibrium system, exhibits periodic changes in reactant concentrations through autocatalysis and feedback regulation. It is governed by nonlinear dynamics, where activators, inhibitors, and intermediates interact to produce sustained oscillations (18). These features make the BZ reaction an ideal theoretical model to study oscillatory phenomena.

A particularly compelling biological system that mirrors the dynamics of the BZ reaction is mitochondrial fission and fusion. Mitochondria, the energy hubs of the cell, undergo constant morphological changes regulated by a balance between fission and fusion processes (19,20). Fission, mediated by DRP1 and associated proteins (21–23), promotes mitochondrial fragmentation, whereas fusion, driven by MFN1/2 and OPA1 (24–26), facilitates mitochondrial elongation and connectivity. These opposing processes are tightly controlled, with molecular factors exhibiting periodic activation and inhibition to maintain mitochondrial homeostasis.

In this study, we explored mitochondrial fission and fusion dynamics through the framework of the BZ reaction. By examining the parallels between these two oscillatory systems, we aimed to uncover how nonlinear feedback and periodic regulation underlie mitochondrial morphology changes. The BZ reaction provides a mathematical and conceptual scaffold to investigate the mechanisms driving the cyclic transitions in mitochondrial dynamics. We performed dynamic simulations of the mitochondrial dynamics very similar to the BZ reaction, where we considered the fission factor as the activator, the intermediary factors as the intermediary factor seen in the BZ reaction and the fusion factor as the inhibitors of the process. After establishing the oscillatory nature of the process, we modified the model to mimic diseased (before treatment) and therapeutic (post treatment) states to determine the changes in the dynamics. Here we observed that under diseased state the levels of the fission and fusion factors increased accompanied by a reduction in the levels of the intermediary factor. But upon therapy we observed a flip where the levels of the fission and fusion factors decreased but the levels of the intermediary factor increased. Further, to determine the biological relevance of the observation, we compared the simulation data with publicly available data and observed similar results. This was further supported by patient survival data. Thus, this approach not only deepens our understanding of mitochondrial regulation but also highlights the potential of chemical oscillators as models for complex biological processes.

## Results

### BZ reaction and mitochondrial fission fusion dynamics

The BZ reaction is a canonical example for a chemical oscillator. These oscillators are characterized by sustained and periodic variations in reactant and intermediate concentrations under non-equilibrium conditions (27). The process is best explained by the Field-Körös-Noyes (FKN) mechanism where oxidization of Ce^3+^ to Ce^4+^ by bromate ions (HBrO_3_) is demonstrated through autocatalysis. And in parallel reduction of Ce^4+^ to Ce^3+^ is brought about by bromide ions (Br^-^) that are generated by malonic acid derivatives. These opposing redox reactions results in the emergence of Ce^4+^ oscillations, and these oscillations are sustained as a result of delay in the feedback via bromide production and autocatalysis (**Fig. 1A**) (28).

**Fig. 1:**
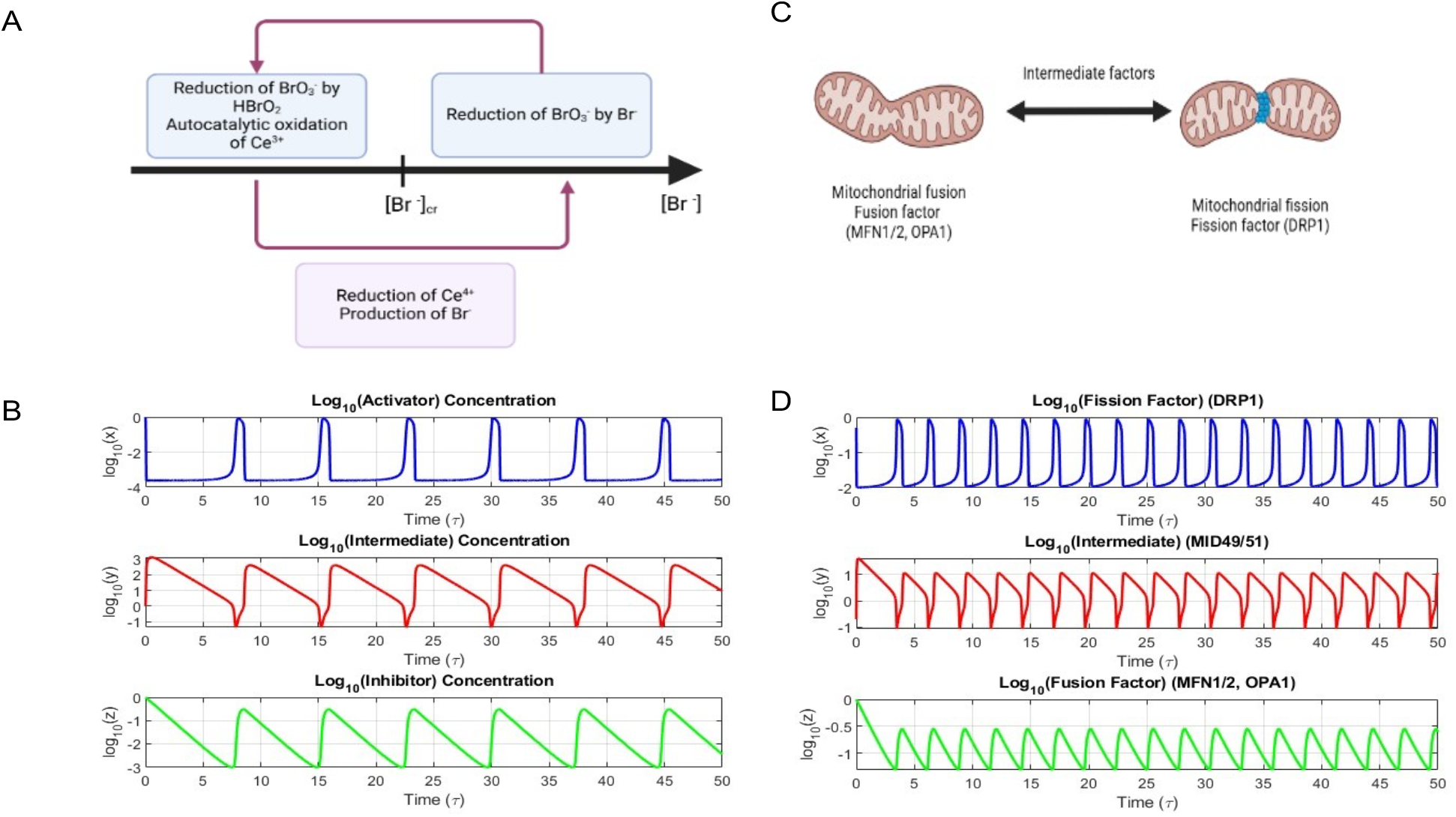
Similarities between BZ reaction and mitochondrial fission and fusion dynamics. **A**) Schematic representation of BZ reaction. **B**) Oscillation profile of activator (blue curve), intermediate (red curve) and inhibitors (green curve) of BZ reaction. **C**) Schematic representation of mitochondrial fission and fusion. **D**) oscillation profile of fission factor (blue curve), intermediate factor (red curve) and fusion factor (green curve).

The oscillations of the BZ reactions can be represented by the simple oregonator model (29). The oregonator model is a simplified FKN mechanism with three variables. Here nonlinear ordinary differential equations (ODEs) are used to define the activator (X, e.g., HBrO2), an inhibitor (Z, e.g., Ce^4+^), and an intermediate (Y, e.g., Br^−^) (**Table 1**). Solution of these ODEs show limit-cycle oscillations (**Fig. 1B**) and hence can be used as an effective minimal framework to understand chemical oscillations.

**Table 1:**
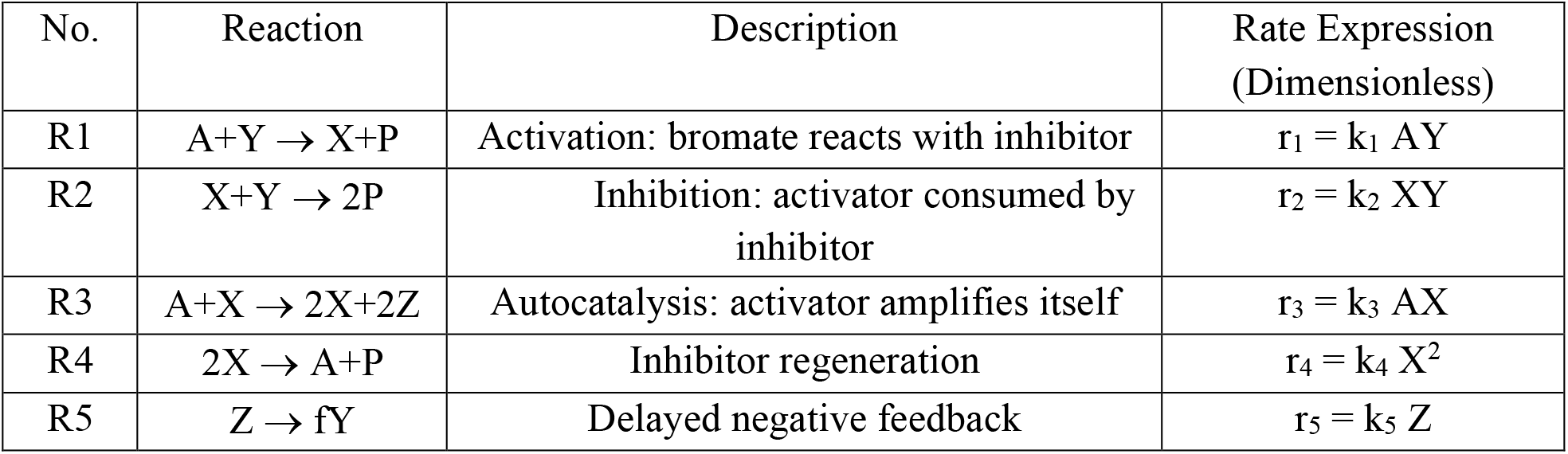
Equations describing the Oregonator model.

| No. | Reaction | Description | Rate Expression (Dimensionless) |
| --- | --- | --- | --- |
| R1 | $A+Y \rightarrow X+P$ | Activation: bromate reacts with inhibitor | $r_1 = k_1 AY$ |
| R2 | $X+Y \rightarrow 2P$ | Inhibition: activator consumed by inhibitor | $r_2 = k_2 XY$ |
| R3 | $A+X \rightarrow 2X+2Z$ | Autocatalysis: activator amplifies itself | $r_3 = k_3 AX$ |
| R4 | $2X \rightarrow A+P$ | Inhibitor regeneration | $r_4 = k_4 X^2$ |
| R5 | $Z \rightarrow fY$ | Delayed negative feedback | $r_5 = k_5 Z$ |

The dynamics of the mitochondrial processes of fission and fusion represent similar oscillatory behavior. The fusion dynamics is controlled by factors like the mitofusins, namely OPA1, MFN1 and MFN2 (24–26). In contrast to this the fission dynamics in controlled by a GTPase, Dynamin-related protein 1 (Drp1) (22). These two processes, though opposing in function, are interdependent and coupled through intracellular signaling. The intermediary factors MiD49 and MiD51 play a major role in the recruitment of the fission factors onto the mitochondrial membrane (30,31). This opposing dynamics of fission and fusion, that helps maintain mitochondrial function and morphology, is achieved by the periodic oscillation in the levels of these mitochondrial factors, suggesting a temporal regulation akin to a biochemical oscillator (**Fig. 1C**). In our model we have aimed to map the mitochondrial regulators onto the Oregonator model variables (**Table 2**) and employed coupled ODEs, similar to that of the Oregonator model, to simulate the time evolution of DRP1, MiD49/51, and MFN1/2–OPA1 (**Table 3**).

**Table 2:** Mapping of variables of the mitochondrial model to the Oregonator model. In this model we observe periodic oscillations in the levels of the fission (DRP1), fusion (MFN1/2, OPA1) and intermediate (MiD49/51) factors (**Fig 1D**) as a result of the coupled feedback loops, similar to that seen in the oregonator model. This model helps us in understanding the dynamics of mitochondrial function and shape as a result of the temporal regulation of two opposing forces similar to that of a BZ reaction where coupled redox reactions maintain periodicity.

| Oregonator model variable | Mitochondrial variable | Function |
| --- | --- | --- |
| Activator (X) | DRP1 | DRP1 initiates and sustains fission activity, analogous to autocatalytic production in the BZ reaction |
| Intermediate (Y) | MiD49/51 | These proteins mediate DRP1 recruitment, temporally linking fusion and fission. |
| Inhibitor (Z) | MFN1/2, OPA1 | Fusion factors counterbalance fission and suppress DRP1 accumulation, acting as negative feedback regulators. |
| A: constant bromate source; X: activator (e.g., $\text{HBrO}_2$); Y: inhibitor (e.g., $\text{Br}^-$); Z: oxidized species (e.g., $\text{Ce}^{4+}$); f: stoichiometric factor in feedback loop; The rate expressions are used to derive the dimensionless Oregonator equations. | | |

**Table 3:** Equations describing the mitochondrial model.

| No. | Biological Interaction | Description | Rate Expression (Dimensionless) |
| --- | --- | --- | --- |
| R1 | $\text{MFN1/2} + \text{DRP1} \rightarrow \text{MFN1/2-inhibited DRP1}$ | Fusion inhibits fission | $r_1 = \epsilon_1 YX$ |
| R2 | $\text{DRP1} + \text{MiD} \rightarrow \text{DRP1 activated (oligomerization)}$ | DRP1 activation via MiD49/51 recruitment | $r_2 = \epsilon_2 XZ$ |
| R3 | $\text{DRP1} + \text{MFN1/2} \rightarrow \text{DRP1 inactivation and MFN degradation}$ | Feedback from fusion to suppress fission | $r_3 = \epsilon_3 XY$ |
| R4 | DRP1 self-activation →<br>active DRP1 (positive<br>feedback) | DRP1<br>oligomerization/autocatalysis | $r_4 = \epsilon_4 X^2$ |
| R5 | MFN1/2/OPA1 synthesis<br>from MiD-regulated<br>feedback | Fusion protein<br>recovery/compensation | $r_5 = fZ$ |
| X: DRP1 (fission activator); Y: MFN1/2 and OPA1 (fusion inhibitors); Z: MiD49/51 (intermediate recruitment regulators); f: Feedback strength (reflecting fusion recovery under high fission); $\epsilon_i$: Corresponding scaled rate parameters | | | |

**Table 4:** Parametric values.

| Sl. No | Parameter | Value Oregonator | Value Normal | Value Diseased | Value Therapy |
| --- | --- | --- | --- | --- | --- |
| 1 | $\epsilon$ | $3.6 \times 10^{-2}$ | 0.01 | 0.005 | 0.005 |
| 2 | $\epsilon'$ | $1.2 \times 10^{-4}$ | 0.001 | 0.0005 | 0.0005 |
| 3 | $q$ | $2.4 \times 10^{-4}$ | 0.01 | 0.02 | 0.02 |
| 4 | $f$ | 1 | 1 | 0.6 | 1.2 |

### Oscillatory behavior of mitochondrial dynamics

Phase plane analysis of the mitochondrial dynamics shows the oscillatory behavior of the process. The phase plane plot of the fission factor (x) and the intermediate factor (y) shows a nonlinear trajectory (**Fig. 2A**). The trajectories converge to stable periodic orbits, highlighting the oscillatory behavior between the fission factor and intermediary factors. The phase plot for the fission factor (x) with the fusion factor (z) demonstrates an antiphase relationship between the factors (**Fig. 2B**). The balance between the fission and fusion dynamics can be traced by periodic cycles.

**Fig. 2:**
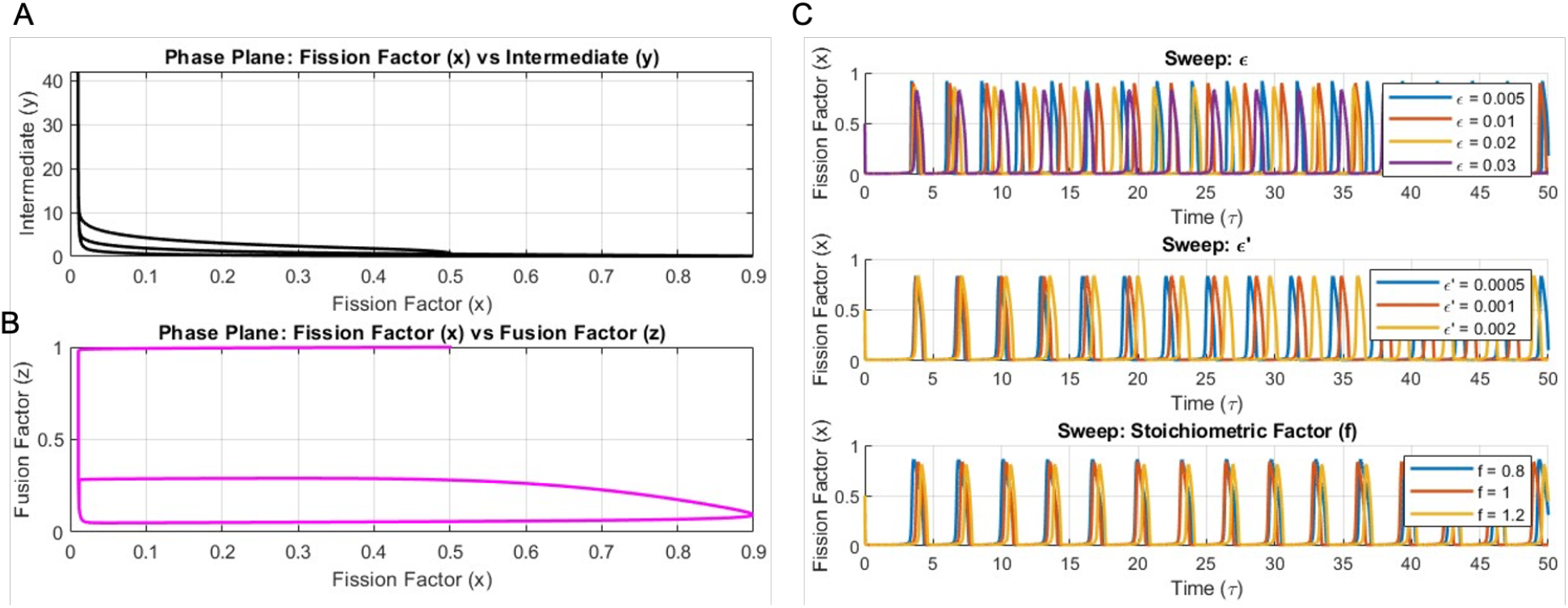
Oscillatory nature of mitochondrial dynamics. **A**) Phase plane plot of intermediate factor vs fission factor. **B**) Phase plane plot of fusion factor vs fission factor. **C**) Parametric sensitivity of *ϵ, ϵ*^′^ and ‘f’.

Next, we performed a parameter sensitivity analysis to determine the parametric effects on the dynamics. The first panel in **Fig 2C** indicates that when the parameter *ε* (indicative of the system’s time scale) is varied there is variation in the oscillatory dynamics. The oscillatory dynamics and *ϵ* values have an inverse relationship; when *ε* decreases there is an increase in the oscillatory frequency. The second panel in **Fig 2C** explores the system sensitivity to the parameter *ϵ*^′^, that influences the intermediary factor dynamics. Similar to *ε*, here we again see an inverse relationship between the parameter and the dynamics, but when *ε*’ decreases, we observe an increase in the frequency of oscillation. But the magnitude of change is smaller than that observed for the variation of the parameter *ε*. The last panel in **Fig 2C** shows the changes in the oscillatory dynamics as a function of changes in the stoichiometric factor that modulates the production rate of the fusion factor (f). Variations in values of ‘f’ alters the amplitude and the periodic nature, highlighting its role in maintaining the strength and stability of the oscillatory cycles. These results highlight the dependence of the system dynamics on the key biochemical parameters.

### Oscillatory dynamics under cancer like condition

To determine the changes in the mitochondrial dynamics under diseased conditions, like that in cancerous cells, we modified our model parameters to mimic increased mitochondrial fission, a characteristic feature of cancer cells. This was achieved by reduction in the stoichiometric factor ‘f’, that determines the production rate and influence of the fusion factor on the system dynamics (**Fig. 3A**). When fission was increased, we observed an increase in the frequency of the fission factor accompanied by a small increase in amplitude but a very large increase in the wavelength (**Fig. 3A**,Top panel)). These observations are consistent with the condition of increased fission. As opposed to the fission factor, under the diseased condition we see a major reduction in the amplitude and wavelength of the intermediary factor accompanied by an increase in the frequency (**Fig. 3A**,Middle panel)). This indicates that the intermediary factors cycle faster under diseased condition to meet the increased need of the cells to undergo fission. But the fusion factor showed no major change in the frequency but a drastic increase in its concentration under diseased condition (**Fig. 3A**,Bottom panel)). Also, when the waveforms are compared, we observe considerable reduction in both amplitude and wavelength indicating the fast cycling of the molecules. We then compared these simulation results with the *in-vitro* results observed in esophageal squamous cell carcinoma (ESCC) upon overexpression of DRP1 (GSE182710) (**Fig. 3B**) (32). Here, we observed that upon overexpression of DRP1 there is an increase in the levels of the fusion factors MFN1 and OPA1 similar to our simulation (Fig 3A,Bottom panel)).

**Fig. 3:**
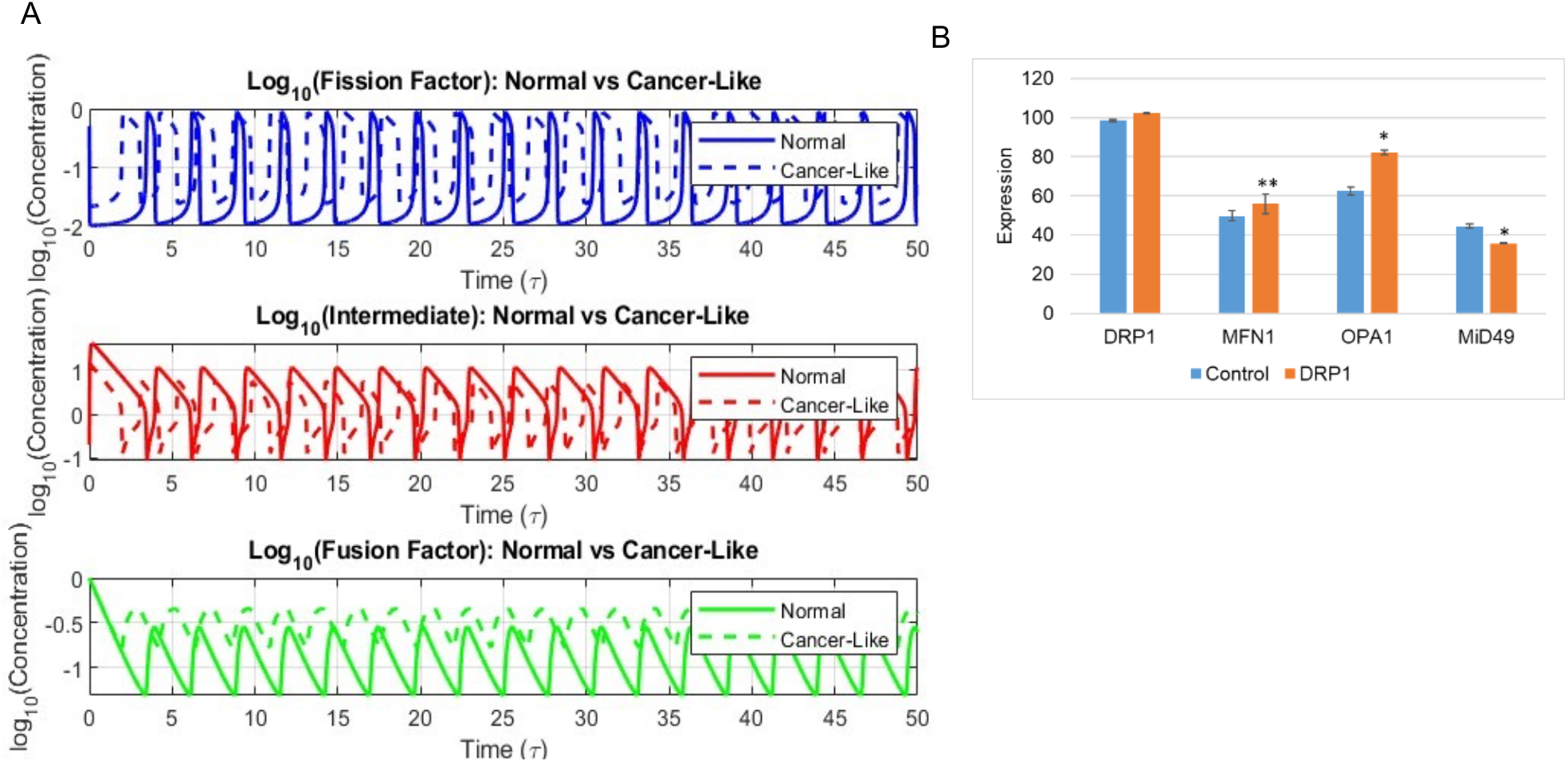
Change in mitochondrial dynamics under cancerous condition. **A**) Comparison of oscillatory dynamics of fission factor (blue curve), intermediate factor (red curve) and fusion factor (green curve) under normal condition (solid curve) and cancerous condition (dotted curve). **B**) Changes in the levels of mitochondrial fission and fusion protein upon DRP1 overexpression in the dataset GSE182710. (* indicates *p-value* <0.01 and ** indicates *p-value* <0.001 calculates using 2-tailed student’s t-test with unequal variance)

Also, consistent with our simulation, we observe a reduction in the levels of the fission factor MiD49 (**Fig. 3A**,Middle panel)). These results indicates that our simulations can effectively mimic and portray the biological conditions.

### Changes in oscillatory dynamics upon therapeutic intervention

Therapeutic intervention that results in the restoration of mitochondrial fusion was simulated by increasing the value of the stoichiometric factor ‘f’. When the stoichiometric factor was increased, we observed a decrease in the frequency of the fission factor accompanied by a very large decrease in the wavelength (**Fig. 4A**,Top panel)). These observations are consistent with the condition of increased fusion. But upon therapeutic intervention we see an increase in the amplitude and wavelength of the intermediary factor accompanied by a negligible increase in the frequency (**Fig. 4A**,Middle panel)). This indicates that the intermediary factors cycle at a slower rate when compared to the diseased condition. Similar to the diseased state, the fusion factor showed no major change in the frequency upon treatment, but a reduction in its concentration was observed (**Fig. 4A**,Bottom panel)). Also, when the waveforms are compared, we observe considerable reduction in both amplitude and wavelength. Here again we compared these simulation results with the *in-vitro* results observed in small cell lung cancer (SCLC) upon the upon treatment with DRP1 inhibitors (GSE267928) (**Fig. 4B**) (33). Here we observed that when the cells are treated with the drugs there is a decrease in the levels of the fusion factors MFN1 and OPA1 similar to our simulation (**Fig. 4A**,Bottom panel)). Also, consistent with our simulation, we observe an increase in the levels of the fission factor MiD51 (**Fig. 4A**,Middle panel)).

**Fig. 4:**
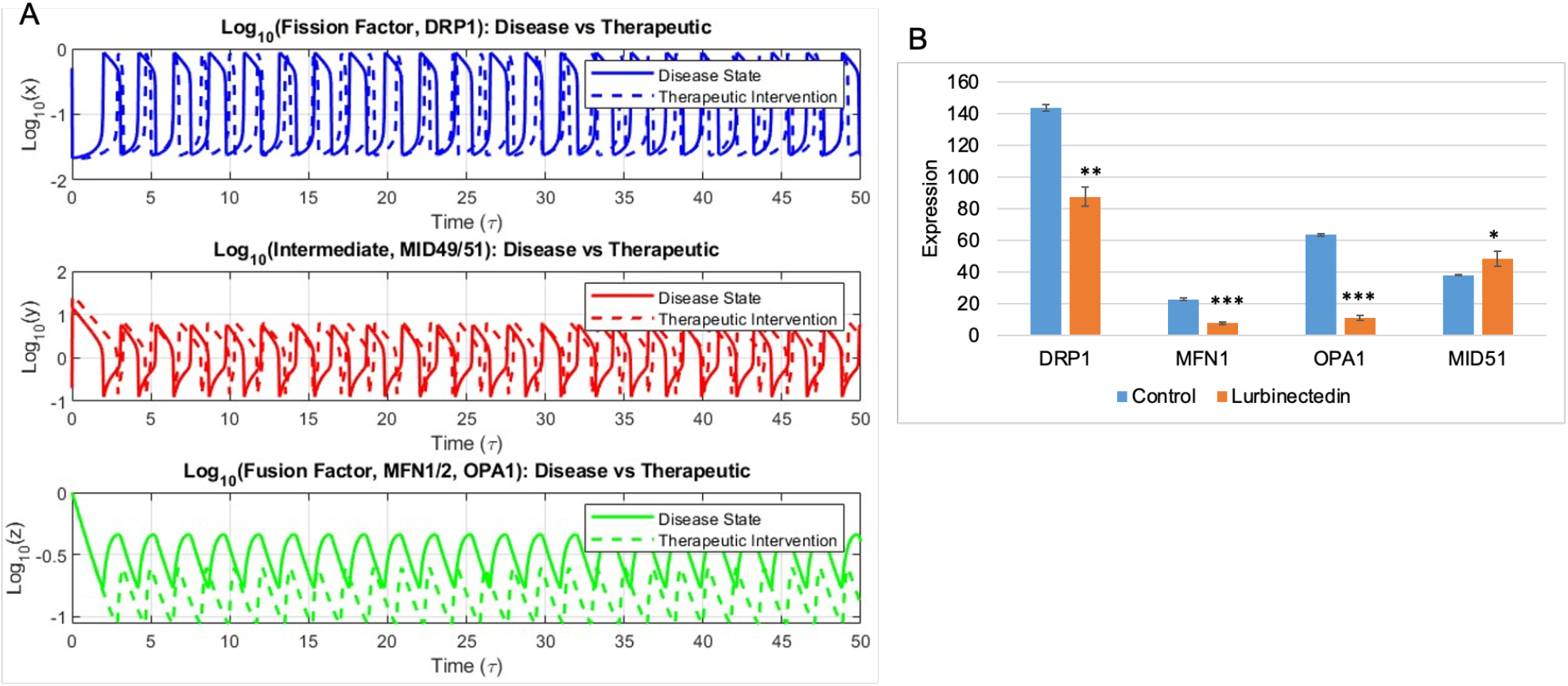
Change in mitochondrial dynamics upon therapeutic intervention. **A**) Comparison of oscillatory dynamics of fission factor (blue curve), intermediate factor (red curve) and fusion factor (green curve) under cancerous condition (solid curve) and upon therapeutic intervention (dotted curve). **B**) Changes in the levels of mitochondrial fission and fusion protein upon treatment with DRP1 inhibitors in the dataset GSE267928. (* indicates *p-value* <0.05, ** indicates *p-value* <0.01 and *** indicates *p-value* <0.0001 calculates using 2-tailed student’s t-test with unequal variance)

### Patient survival and mitochondrial dynamics

Next, we determined the relationship between the different fission, fusion and intermediary factors in cancer. To this end, we interrogated the Gene Expression Omnibus (GEO), a public functional genomics data repository supporting MIAME-compliant data submissions run by the National Center for Biotechnology Information (NCBI) (https://www.ncbi.nlm.nih.gov/geo). We accessed lung cancer lung (GSE30219) (34) and breast cancer patient datasets (GSE45255, GSE11121, GSE20685 and GSE12276) (35–38). We found that in lung cancer patients with low levels of the fission factor DRP1showed better survival (**Fig. 5A**). Similarly, the fusion factors MFN1 (**Fig. 5B**) and OPA1 (**Fig. 5C**) showed a negative correlation with patient survival where high expression of the factors resulted in poor patient survival. But as opposed to the fission and fusion factors the intermediary factor MiD49 showed a positive correlation with patient survival where high levels of the protein resulted in better patient survival (**Fig. 5D**). We saw similar trends when we analyzed patient survival in breast cancer datasets based on the levels of the mitochondrial dynamic’s proteins. Patients showed poor survival when they had elevated levels of the fission factor DRP1 (**Fig. 5E**). And when the levels of the fusion factors MFN1 (**Fig. 5F**) and OPA1 (**Fig. 5G**) were high patients again showed poor survival. However, patients with elevated levels of the intermediary factor MiD49 (**Fig. 5H**) showed better survival.

**Fig. 5:**
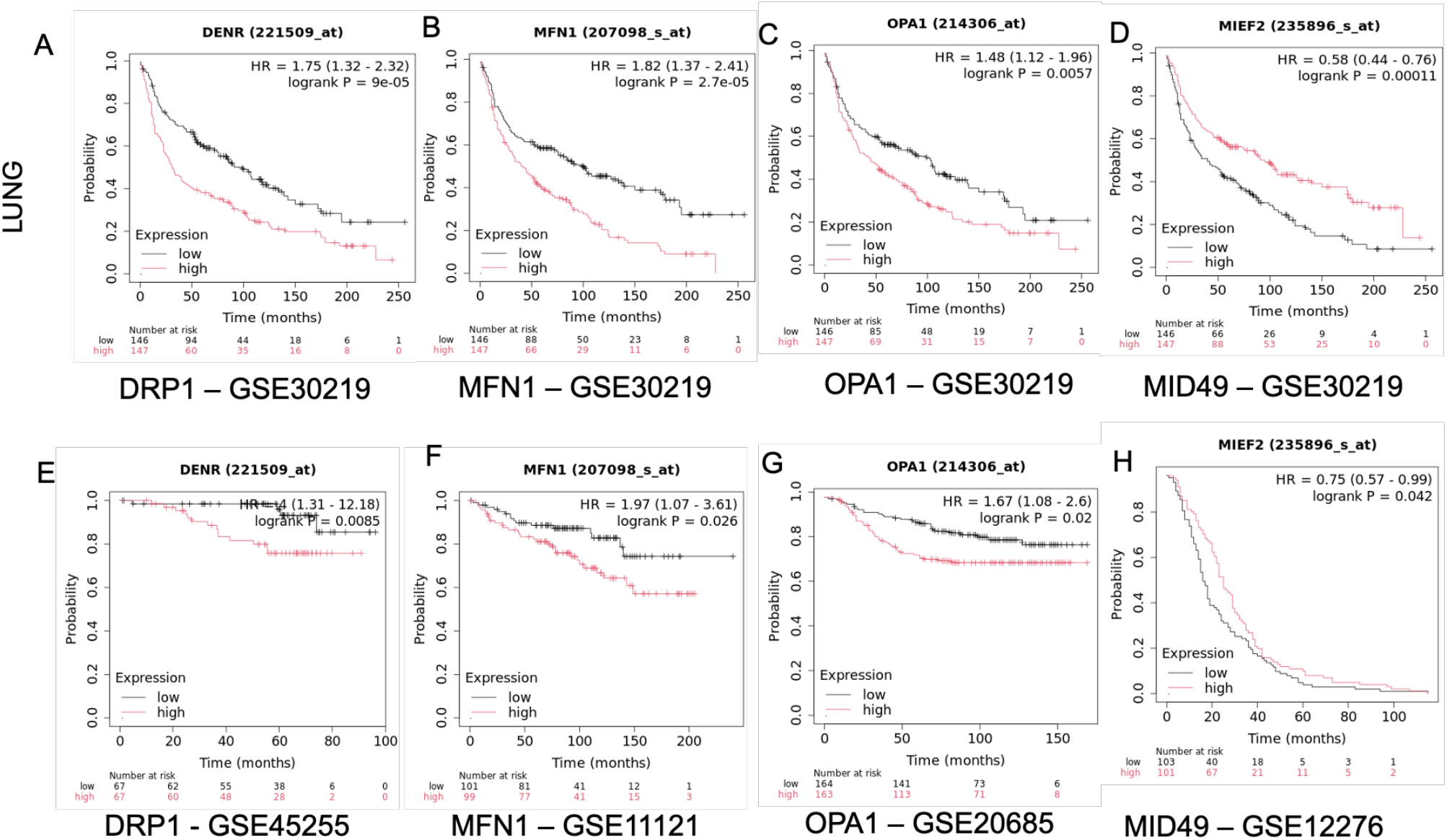
Overall patient survival in relation to the levels of mitochondrial fission and fusion proteins. **A-D**) Survival trends in the lung cancer dataset GSE30219 based on the levels of DRP1, MFN1, OPA1 and MiD49 respectively. **E-H**) Survival trends in breast cancer patients based on the levels of DRP1 (GSE45255), MFN1 (GSE11121), OPA1 (GSE20685) and MiD49 (GSE12276) respectively.

## Discussion

Several theoretical studies aimed at a spatiotemporal understanding of biological control systems regulating biochemical clocks, cellular decision-making, and biological signaling networks, have been reported in the literature. However, only a few have leveraged the power of the BZ system (39). As far as we are aware, this is the first attempt to explore the parallels between mitochondrial fission/fusion dynamics and non-linear chemical oscillations modeled by the BZ reaction. Our results highlight the dynamic nature of mitochondrial fission and fusion processes, emphasizing their cyclic interplay and regulation under normal, pathological (cancer), and therapeutic conditions. Phase plots and the temporal dynamics data show a tightly regulated balance between the processes of mitochondrial fission and fusion which are opposing and yet complementary processes that are essential for the normal cellular functions. In pathological states, we observed an increase in both fission and fusion activity simultaneously, which indicates a mitochondrial hyperactivity that is essential for the metabolic and proliferative demands of cancer cells. Conversely, the intermediary factor, which functions as a transitional regulator of the mitochondrial dynamics network, was significantly reduced in cancer. -However, following treatment, there was a notable increase in the intermediary factor, suggesting a potential restoration of the regulatory balance within the mitochondrial network.

The increased fission and fusion observed under cancer-like conditions reflects a state of increased mitochondrial turnover, often referred to as mitochondrial dynamism (40–42). This hyperactivity enables rapid remodeling of the mitochondrial network, facilitating the cancer cells to meet elevated energy demands and maintain resistance to stress-induced apoptosis (43). Enhanced fission contributes to mitochondrial fragmentation, which is frequently associated with increased glycolytic activity, a hallmark of cancer metabolism (44–46). Concurrently, elevated fusion helps to maintain mitochondrial functionality and mitigate damage by facilitating the exchange of mitochondrial content (47,48). However, the observed reduction in the intermediary factor suggests that despite the hyperactive dynamics, cancer cells may experience a bottleneck in the regulatory processes that mediate the transition between fission and fusion.

Therapeutic interventions significantly reduced both fission and fusion activities, indicating a shift toward a more stable and less dynamic mitochondrial state. This stabilization likely reflects a restoration of mitochondrial homeostasis, improving cellular energy efficiency and reducing oxidative stress (49,50). Notably, the increase in the intermediary factor following treatment points to a recovery of the regulatory processes that balance mitochondrial fission and fusion. This recovery may play a critical role in restoring mitochondrial network functionality, potentially making cells more responsive to apoptotic signals and less adaptable to metabolic stress.

The clinical relevance of these findings is underscored by the observation that mitochondrial dynamics regulators, such as DRP1, MFN1, and OPA1, are tightly linked to the cyclic nature of fission and fusion processes. Alterations in these dynamics offer promising avenues for therapeutic intervention. The ability to modulate mitochondrial fission and fusion provides a unique opportunity to disrupt the adaptive advantages conferred by heightened mitochondrial dynamics in cancer cells.

In summary, this study adapting the principle underlying the BZ reaction, elucidates the dual role of fission and fusion in cancer progression and highlights the therapeutic potential of restoring balance to mitochondrial dynamics. The observed changes in the intermediary factor provide new insights into the regulatory mechanisms of mitochondrial turnover, suggesting that targeting this aspect of mitochondrial dynamics could be a promising strategy for future research and therapeutic development.

## Methods

### Mathematical model

A system of ordinary differential equations was used to define the mitochondrial fission and fusion dynamics. This system of equations was modelled based on the Oregonator model of the Belousov-Zhabotinsky (BZ) reaction. The system of equations described below helps capture the oscillatory nature of mitochondrial fission, fusion and intermediary factors.

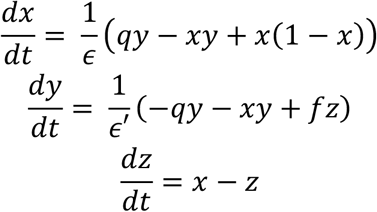

Where,

x = Concentration of fission factor (DRP1)

y = Concentration of intermediary factor (MiD49/51) z = Concentration of fusion factor (MFN1/2, OPA1)

*ϵ, ϵ*^′^, *q* & *f* = System parameters

### Parameter values

The parametric values for the simulation were estimated and scaled based on the original Field-Noyes equations for the BZ reaction and biologically relevant stoichiometric relationships observed in mitochondrial dynamics. The parametric values used for the simulation are as follows.

### Software and dataset

The computational analysis of the mathematical model was simulated using MATLAB (version 2023b). The codes used for the simulation are available on GitHub (https://github.com/ARShuba/Mitochondrial_Dynamics). The datasets were downloaded using *GEOquery* R Bioconductor package (51). The data was processed to obtain gene wise expression using R (version 4.0.0).

### Kaplan-Meier analysis

KM-plotter (52,53) was used for Kaplan-Meir analysis. The data was separated based on the median levels of gene expression. The number of samples in protein high and low categories are given below.

GSE30219 (Lung cancer) – DRP1 – n(High) = 147, n(Low) = 146

GSE30219 (Lung cancer) – MFN1 – n(High) = 147, n(Low) = 146

GSE30219 (Lung cancer) –OPA1 – n(High) = 147, n(Low) = 146

GSE30219 (Lung cancer) – MiD49 – n(High) = 147, n(Low) = 146

GSE45255 (Breast cancer) – DRP1 – n(High) = 67, n(Low) = 67

GSE11121 (Breast cancer) – MFN1 – n(High) = 99, n(Low) = 101

GSE20685 (Breast cancer) – OPA1 – n(High) = 163, n(Low) = 164

GSE12276 (Breast cancer) – MiD49 – n(High) = 101, n(Low) = 103

## Data Availability

The datasets used in the study are available in online repositories as mentioned in the methods. The codes used for the analysis are available on GitHub (https://github.com/ARShuba/Mitochondrial_Dynamics).

